# Temporal GWAS identifies a widely distributed novel protein Stv contributing to pathogen success in *Shigella* spp

**DOI:** 10.1101/2022.08.23.504947

**Authors:** P. Malaka De Silva, Rebecca J Bennet, Francesco Costa, Antonina Andreeva, Rebecca J Bengtsson, Malcolm J Horsburgh, Tim R Blower, Alex Bateman, Kate S Baker

## Abstract

*Shigella* has emerged as a successful pathogen posing a threat to human health worldwide in the recent years. The rise of *Shigella* over the years has variably been attributed to AMR genes, virulence, and bacterial competition factor but no method has taken a function-agnostic approach to look for factors associated with modern variants of the pathogens. To address this gap, here we combined historical and modern isolate collections to identify such factors through a novel approach, termed temporal GWAS (tGWAS). Our analyses identified a novel putative adhesin gene, which we called *stv* that was associated with time of isolation and concentrated in expanding lineages of *Shigella* spp, as well as widely distributed in other bacterial species. We confirmed that *stv* is carried on a small 2689bp plasmid and *in silico* analyses revealed that Stv contained a new protein domain that was in combination with other domains in a manner suggestive of a secreted bacterial toxin. However, an all-proteome AlphaFold interaction screen indicated a high likelihood of interactions with fimbrial proteins, despite fimbrae thought to be defunct in *Shigella* sp. Collectively, these findings suggest that Stv is a novel protein domain with a likely role in *Shigella* success that is also widely distributed in other species.

## Introduction

*Shigella* spp. are the causative agent of shigellosis with the main symptoms being diarrhoea (often bloody), abdominal pain, stomach cramps, and fever (1–3). Although the genus is comprised of four species, most (86-90%) of the global disease burden is caused between *S. flexneri* and *S. sonnei* (4,5). Children under the age of five years old from lower- to middle-income countries are disproportionately affected mainly due to the poor water sanitation and hygiene where *S. flexneri* is endemic (6,7) while shigellosis in high income countries are often attributed to travel to endemic areas and close-contact communities is dominated by *S. sonnei* (8–12).

The success of *Shigella* as pathogens is partly attributable to its ability to adapt and evolve efficiently (13). Antimicrobial resistance (AMR) has also been an exacerbating factor, with *Shigella* now being WHO AMR priority pathogens (14). These AMR determinants are often associated with mobile genetic elements (MGEs) facilitating horizontal gene transfer and contribute to the accessory genome of *Shigella* spp. (15). *Shigella* has also streamlined its arsenal of virulence factors and incorporated other useful proteins such as colicins for interbacterial competition (16). It has also undergone genome reduction to dispose of immunogenic components such as some of the fimbriae genes, which are unfavourable from an immune evasion perspective (17). Hence, the evolution of *Shigella* as a pathogen over the years appears to have been multifactorial, but to date no agnostic screen of which accessory genome determinants are changing over time has been completed.

Historical isolate collections provide an invaluable resource to look at pathogen evolution (18). One such historical collection is the Murray collection, consisting of several hundred *Enterobacteriaceae* collected from a wide range of geographic areas in the pre-antibiotic era (1917 – 1954) and curated by the eminent microbiologist, Professor Everitt George Dunne Murray (19). It features ∼90 *Shigella* spp., and the whole collection, which is held by the National Collection of Type Cultures (NCTC), has been genome sequenced as a public resource for scientific research (13). The immense utility of the Murray collection has already been demonstrated via seminal studies elucidating the profile of pre-antibiotic era *Enterobacteriaceae* plasmids, the evolution of key phenotypes in *Klebsiella* spp., and the evolution of antimicrobial resistance (20–22).

Here, we aimed to identify those accessory genome changes that are associated with modern *Shigella* spp. to characterise factors that have contributed to the long-term success of the pathogen. To do so, we combined the Murray collection genomes with those from more contemporary *Shigella* isolates to create collections spanning the antibiotic era. We describe our novel strategy as a temporal Genome Wide Association Study (tGWAS), where genetic factors positively associated with time (as a continuous variable) are identified as putative contributors to pathogen success. The efficacy of our approach is demonstrated by the recovery of AMR genes, which are known to have increased over this time frame, virulence factors, and the discovery of novel uncharacterised determinants that contribute to pathogen success in *Shigella* spp. and are widely distributed in other pathogenic organisms.

## Results and discussion

### tGWAS identifies novel genetic features associated with success in S. flexneri

To assemble an appropriate dataset for interrogating *Shigella* genome evolution over the antibiotic era, we first examined the context of the *S. flexneri* in the Murray collection (n=45) alongside a more contemporary yet temporally and geographically diverse *S. flexneri* collection (n=262) (23). Our phylogenetic reconstruction (see methods) using 84,391 single nucleotide polymorphisms (SNPs) from core regions revealed that Murray collection isolates clustered with Phylogroup (PG) 1 through PG5 (n=20 for PG1, n=3 for PG2, n=14 for PG3, n=6 for PG4, n=2 for PG5) but not with PG6 or PG7 (Figure 1B) indicating both co-existence and clonal replacement of *S. flexneri* PGs, which agrees with previous studies (23) (Figure 1A). This showed that the Murray collection isolates represent an appropriately diverse retrospective collection of *S. flexneri* on which to conduct gene discovery.

**Figure 1.**
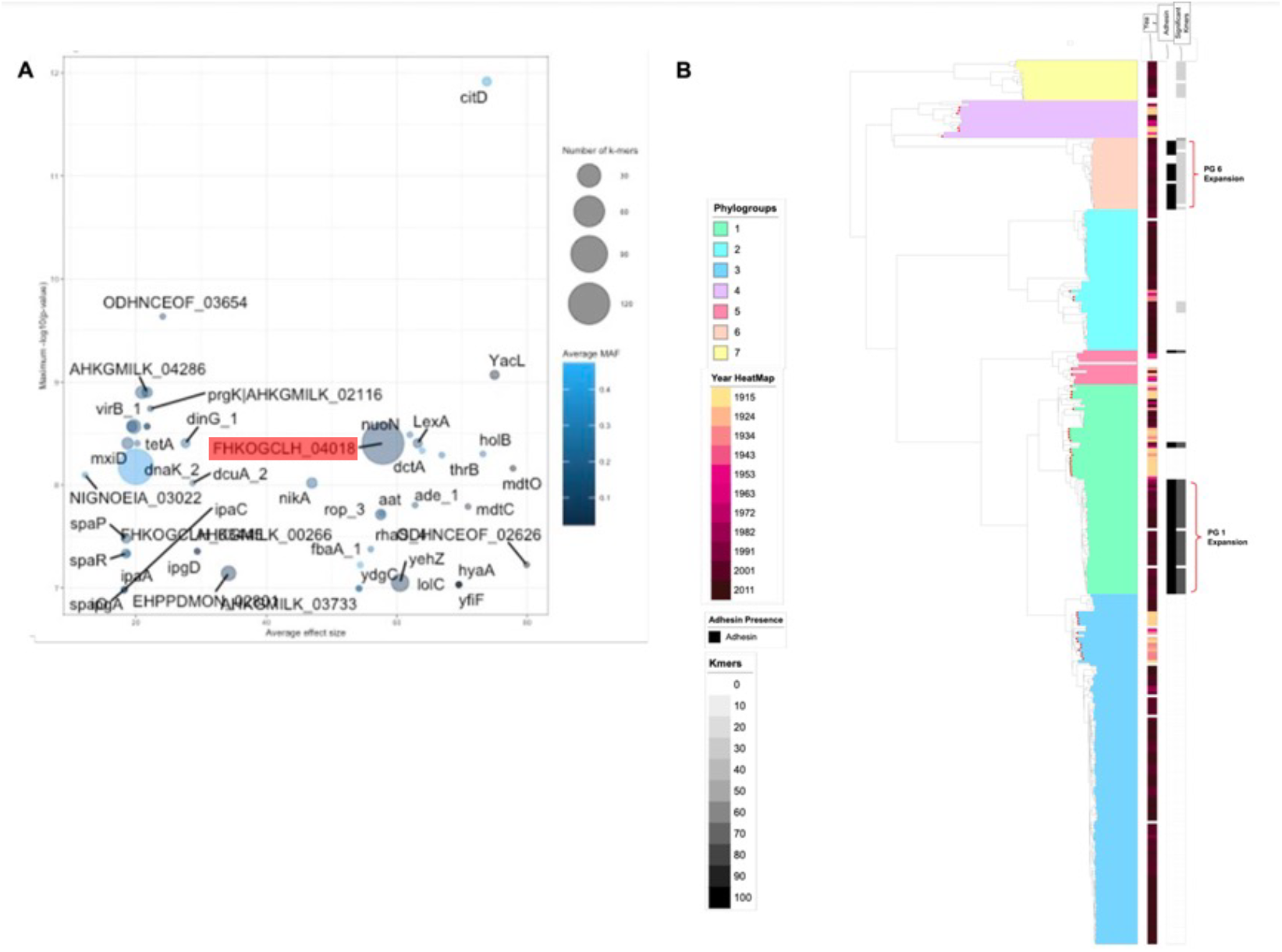
*stv* is a novel protein associated with clonal expansions in contemporary *S. flexneri*. (A) The bubble plot shows tGWAS results for kmers (n=306) by corresponding gene. Bubbles scaled by the number of associated kmers (bubble size) and shaded by the average minor allele frequency (MAF) are displayed as a scatter plot by average effect size (x-axis) and statistical support (negative logarithm of the *p*-value, y-axis). In this plot, *stv* is labelled as FHKOGCLH_04018 and is highlighted in red. (B) A mid-point rooted maximum likelihood phylogeny of *S. flexneri* isolates (n=309) is shown with red dots overlying tips derived from Murray collection isolates. Phylogroups (PGs) are shown shaded over the phylogeny and coloured according to the inlaid key. Isolate year of isolation is shown in the leftmost metadata track, coloured by inlaid heatmap. The binary presence of *stv* is shown as a middle track with black indicating presence and white indicating absence, and the number of significant *stv-*associated kmers in each isolate is shown by a grey scale heatmap in the rightmost track. Two *stv-*associated phylogroup expansion are indicated by labelled brackets

To confirm our suspicion that AMR genes could act as a positive control when looking for factors increasing over time, we determined whether AMR genes had increased over time. Specifically, Murray collection isolates had a lower number of AMR determinants per isolate (n=45, 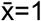) than modern isolates (n=246, 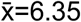) (T test: t_291_ = 11.6, *p*<0.001) (Supplementary Table 1). This is consistent with previously described trends in *S. dysenteriae* Type 1 (24) and other bacterial pathogens (22). We also observed a higher number of virulence genes per isolate among the modern isolates (n=246, 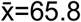) compared with those in the Murray collection (n=45, 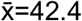) (t_291_ = 7.15, *p*<0.001, Supplementary Table 1) indicating an increase in both AMR and virulence genes over time, consistent with previous studies (20,25).

To identify novel factors that were found to be increasing in *S. flexneri* populations over time, we then conducted our novel approach temporal GWAS. To do so, we conducted bacterial GWAS to identify genetic variation (i.e. SNPs, clusters of orthologous groups [COGs], and kmers) that that was positively associated with time (in year, as a continuous variable). This identified genetic variants with significant associations with time across SNPs (n=93, minlog_10_(*p*)>2), COGs (n=359, lrt *p*<0.05), and kmers (n=306, *p*<1.04e-07) (Supplementary Table 2). We focused on the latter group and found kmers (between 10 and 100bp in length) were spread through 47 coding sequences where 85% (n=40/47) were contained within coding sequences of proteins of known function (Figure 1B). These reflected our findings of increasing AMR and virulence in archived *S. flexneri* as we recovered kmers relating to known AMR genes *mdtC, mtdO* and *tetA* (26,27) and different components of the Type 3 Secretion System (T3SS) including the assembly machinery (*spaO/P/R*), effector proteins and their chaperons (*ipaA/C, icsB, ipgA*), and regulators (*mxiC, virB*) (28,29). This demonstrated the efficacy of our tGWAS approach for identifying determinants contributing to pathogen success over time.

Critically however, our tGWAS also recovered kmers relating to genes encoding proteins of unknown function (15%, n= 7/47 genes) (Figure 1B). In fact, the gene with the highest overall number of kmers mapping (n=123) was one of unknown function with high effect size and statistical significance scores (FHKOGCLH_04018, Figure 1B). The highest BLASTp match for this coding sequence (with a 99% amino acid sequence identity) was to a putative *E. coli* protein WP_205849698.1, to which there was also good three-dimensional similarity (Supplementary Figure 1). Although this *E. coli* protein was labelled as an adhesin, this appeared to be unsupported by functional characterisation and appeared to have transferred over from a *Bordatella* protein. We termed this novel protein (FHKOGCLH_04018) *stv* and examined its distribution across the *S. flexneri* population structure. This revealed that *stv* variants were polyphyletic across multiple phylogroups and were associated with population expansions in Phylogroups 1 and 6 (Figure 1A), consistent with an accessory genome component contributing to pathogen success over time.

### stv associated with success in S. sonnei and is plasmid encoded

To validate the suggestion that *stv* was associated with epidemiologically successful *Shigella* lineages, we examined the distribution of *stv* across another pan-antibiotic era collection of the next most burdensome *Shigella* species, *S. sonnei* (n=305, Supplementary Table 3 and Methods). This revealed that while *stv* was only sporadically distributed in Lineages 1, 2 and 5, it was acquired and stabilised early in the emergence of Lineage 3; the contemporarily globally dominant Lineage (Figure 2A, (2)). Success of Lineage 3 has been previously attributed to the acquisition of key AMR determinants, particularly, the Tn7/Int2 MDR determinants (30). Our analysis however indicates that *stv* was present in the clonal expansion of into Lineage 3 prior to the acquisition of the Tn7/Int2 cassette (with the first observation of *stv* in Lineage 3 being in 1943 and the Tn7/Int2 cassette being first observed in 1990). Consistent with the findings in *S. flexneri* above, this highlights role *stv* may have played in the success of Lineage 3.

**Figure 2.**
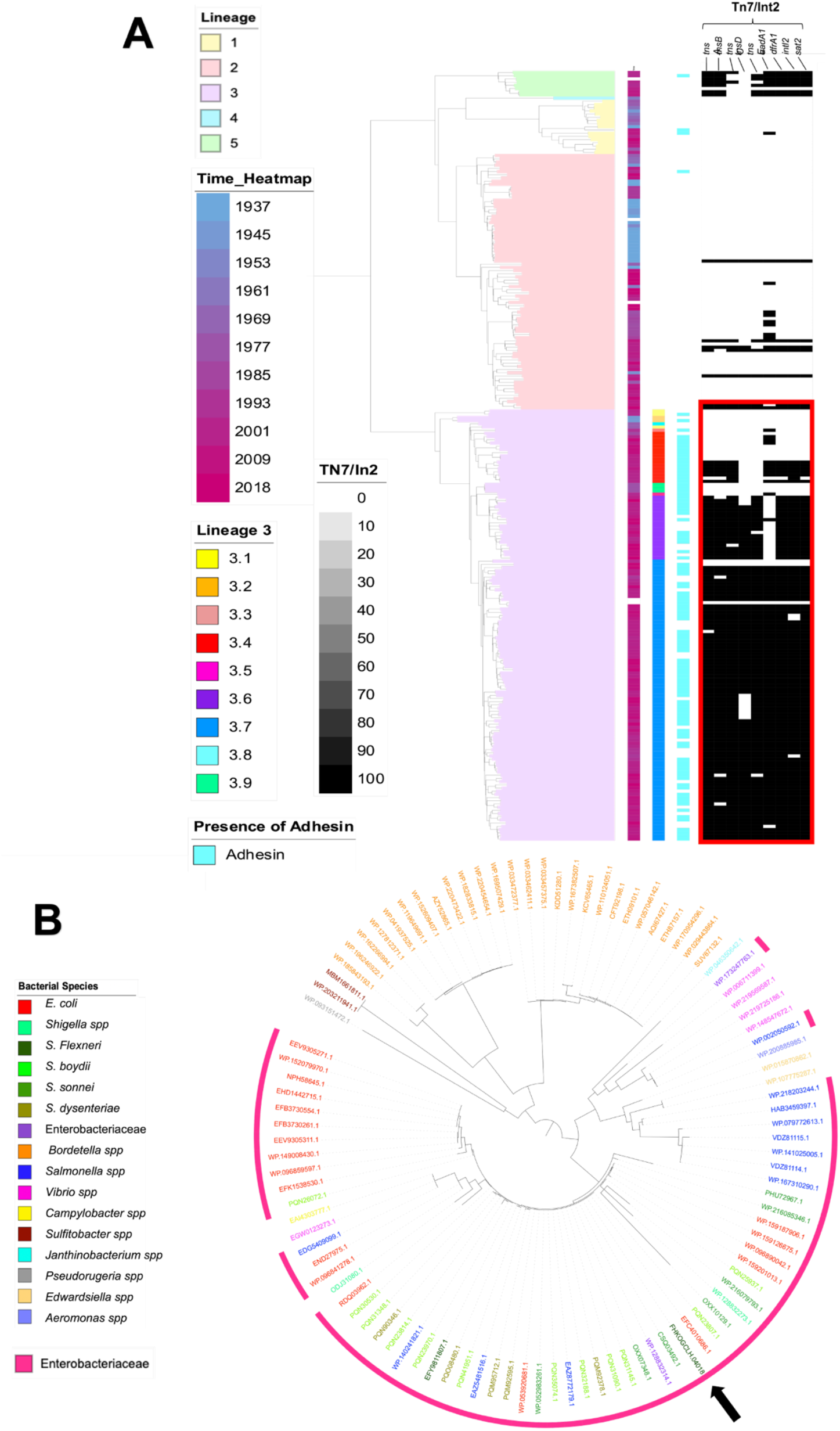
The natural distribution of Stv in other bacterial species. (A) Mid-point rooted phylogeny of 305 *S. sonnei* isolates with the coloured blocks in the phylogeny indicating the lineages of the isolates. First three coloured strips next to the phylogeny represents the isolation year, Lineage 3 genotypes and the presence/absence of *stv* in the isolates from left to right. The rightmost black and white strips with the labelled headings indicate the presence (black) and absence (white) of AMR genes associated with the Tn*7*/Int2 cassette. (B) Phylogeny of the aligned protein sequences most closely related and clustered to Stv and the colours represent the different bacterial species for the protein sequence as indicated by the inlaid key and bacterial species belonging to *Enterobacteriaceae* are indicated with the pink strip around the phylogenetic tree while the black arrow points to the location of the Stv sequence from *S. flexneri* used as the reference.

We examined the genetic context for the *stv* protein and found that *stv* is located on a small plasmid. Specifically, a contiguous sequence of 2689bp was confirmed using bidirectional coverage PCR (Supplementary Figure 2). We named this plasmid pSTV and it contained two other coding sequences; one for a plasmid mobilisation protein (*mobA*) (31) and another for a *rop* family protein *rop3*, which is known to regulate plasmid copy number (32). The other features in pSTV included an oriC type origin of replication and an origin of transfer (oriT). Notably, no AMR genes or addiction systems that indicate selection and/or maintenance in bacterial populations were found in pSTV. Furthermore, estimated copy number for pSTV from mapping the sequencing reads was determined to be ∼9 copies (see methods).

Given that *stv* was carried on a plasmid and showed evidence of convergent acquisition (i.e. was poly-phyletic in our *Shigella* phylogenies), we explored the abundance of *stv* in other bacterial species. Specifically, we found clustered relatives of Stv in the nr database across multiple other bacterial species including other *Enterobacteriaceae* (i.e. *E. coli* and *Salmonella* spp.) and more distantly related species (i.e. *Bordetella* spp.) indicating the widespread nature of Stv and its possible importance of shaping population structures in other pathogen groups (Figure 2B).

### Stv contains a new protein domain found in a variety of protein domain contexts and has structural features of a secreted bacterial toxin

We then sought to uncover the function of Stv protein and since the annotation of protein WP_205849698.1 from *E. coli*, which was the top BLASTp hit for Stv, as an adhesin was not functionally validated. We performed iterative searches within the UniProt Reference Proteome set and identified ∼150 proteins with the same domain. We then added this domain to the Pfam database under the accession PF21527 and explored its distribution. The domain was predominantly found in *Actinobacteria* and *Proteobacteria* (alpha, beta, gamma and delta) and more sporadically in a wider range of bacteria (Supplementary Figure 3).

In further efforts to determine possible functions of Stv, we investigated the other domains associated with the Stv domain (i.e. PF21527) and found a range of different architectures. Notably however, the Stv domain was only observed at the C-terminus of the protein sequences (Figure 3A and Supplementary Figure 3). Perhaps the most intriguing domain architecture we found was the Stv domain present at the C-terminus of proteins containing Rhs repeats (Figure 3A), which are known to be secreted toxins (33). The toxin domain is found at the C-terminus of these proteins and the Rhs repeats form a shell that encapsidates the toxin (PMID:23913273). This strongly suggests that the Stv domain has toxin activity. Some of the other domains we found associated with Stv were also indicative of potential toxin activity, i.e. DUF6861 is an N-terminal integral membrane protein domain which contains a Glycine zipper motif (GXXXGXXXG) that is associated with several other toxin domains from toxin-antitoxin systems (34) such as novel toxin 15 (PF15604), tuberculosis necrotizing toxin TNT:PF14021, Bacterial EndoU nuclease (PF14436) and CdiA C-terminal tRNase domain (PF18664). The Stv domain is also found closely associated with Peptidase C80 domains. Interestingly there are two to four copies of the Peptidase C80/Stv domain pair found in various homologues. As noted by the MEROPS database, Peptidase family C80 contains self-cleaving proteins that are precursors of bacterial toxins. Presumably the autolytic cleavage by the C80 peptidase releases the toxic Stv domain once within a target cell in an analogous way to the RTX toxin.

**Figure 3.**
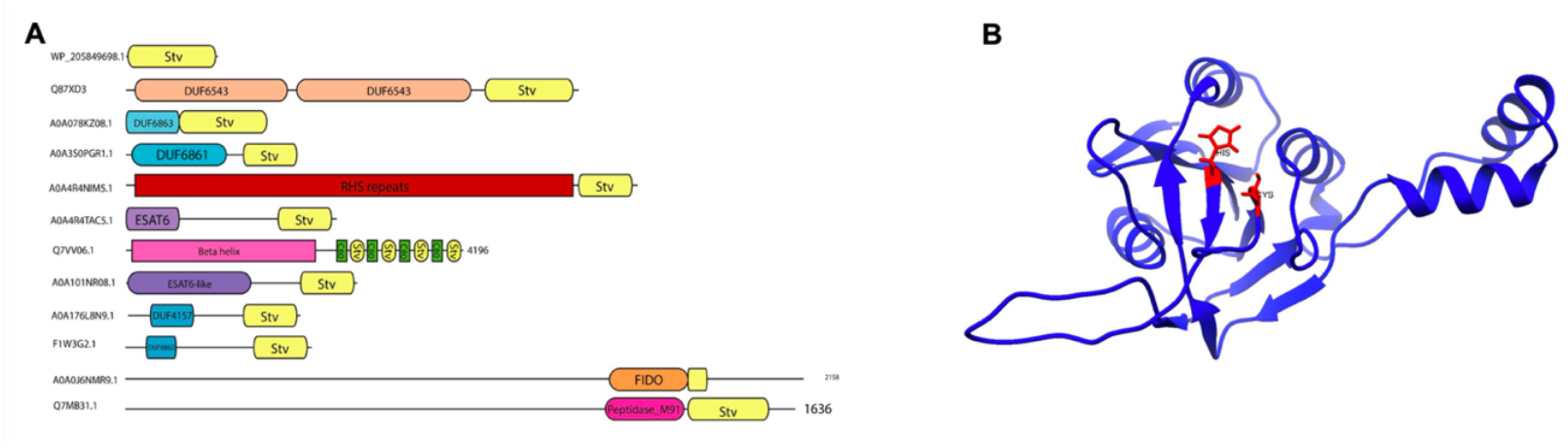
Representative domain architecture of Stv domain containing proteins and structure model of Stv. (A) Domain architectures of representative proteins containing the Stv domain (yellow) with the accession numbers given on the left and where applicable the length of the proteins are given on the right. (B) Structure model of Stv as generated by AlphaFold2 with the conserved histidine and cysteine residues highlighted in.

To further support this, an AlphaFold 2 model of Stv domain was predicted. We conducted a search for homologues using the DALI server to understand the similarity of Stv domain to any other known structures and the top DALI matches are shown in Supplementary Figure 4. However, none of the matches had a high DALI score to indicate a homologous relationship. The predicted structure of the Stv domain was to contain a central mixed ß-sheet flanked on both sides by α-helices and we observed an arrangement of the conserved regions in the Pfam seed alignment of a histidine (His) and a Cysteine (Cys) residue reminiscent of His-Cys diads observed in a number of caspases (35) and bacterial caspase homologues (36), particularly interesting is the similarity to cysteine protease domain (CPD) of MARTX toxins (37) (PDB entries 3fzy, 3pa8, 7d5y, etc) further pointing us towards a Stv being a secreted toxin but perhaps with a proteolytic activity similar to RTX cysteine proteases.

### AlphaFold interaction screening suggests interactions of Stv with fimbrial proteins and haem related functions for virulence

To further explore the potential functions of the Stv protein, we carried out a genome wide AlphaFold screen co-folding Stv with each protein from the predicted protein from the *Shigella* proteome (from a Lineage 3 Genotype 3.7.30.1 strain 627346 from (17)). The top hit was fimbrial-like protein FimF with an ipTM value of 0.858 (Table 1, Supplementary Table 4). This along with another fimbrial protein, FimA, with an ipTM value of 0.732 suggested that there could be interactions with fimbrial proteins (Table 1, Supplementary Table 4).

**Table 1.** Hits with ipTM score > 0.7 from the AlphaFold interaction screening.

| ipTM | UniProt accession | UniProt function |
| --- | --- | --- |
| 0.858 | A0A0H8E7B2 | FimF Fimbrial-like protein |
| 0.837 | N/A | Leader peptide [*E. coli* HB101] |
| 0.824 | A0A0I0NH38 | glutathione-specific gamma-glutamylcyclotransferase |
| 0.820 | A0A0I5CAT7 | NrfF-Formate-dependent nitrite reductase complex subunit - Possible subunit of a haem lyase |
| 0.810 | A0A0H8GMS2 | BioA-Adenosylmethionine-8-amino-7-oxononanoate aminotransferase |
| 0.809 | A0A0I3FMB2 | Peptidyl-dipeptidase DCP |
| 0.808 | A0A0H7Q5S2 | pyrimidine/purine nucleoside phosphorylase |
| 0.804 | A0A0I0XM66 | Sodium/proline symporter |
| 0.797 | A0A0H9P931 | TolR-Part of the Tol-Pal system |
| 0.795 | A0A0H9PH53 | DUF4434 - Tim barrel. Probable enzyme |
| 0.793 | A0A0H7HUR1 | CobU Bifunctional adenosylcobalamin biosynthesis protein |
| 0.791 | A0A0H8FZE5 | 2-dehydro-3-deoxy-phosphogluconate aldolase. |
| 0.790 | A0A0H8MQD9 | FAD assembly factor SdhE |
| 0.789 | A0A0H8LXU7 | Dimethylsulfoxide reductase subunit B |
| 0.783 | A0A0H8LXU7 | Dimethylsulfoxide reductase subunit B |
| 0.779 | A0A0I3RVH3 | Superoxide dismutase [Cu-Zn] |
| 0.778 | A0A0I2A389 | YaeQ family protein |
| 0.775 | A0A0I1BS92 | Uncharacterized plasmid protein |
| 0.770 | A0A8B4CR98 | RecE-Exonuclease family protein |
| 0.762 | A0A0H8TEF8 | DppC Dipeptide transporter |
| 0.733 | A0A0H9WQY6 | Ethanolamine utilization protein EutP |
| 0.731 | A0A0I3QNF6 | Phage Tail protein |
| 0.731 | A0A0I0JV29 | Penicillin G acylase |
| 0.731 | A0A0H9RMA1 | 3-oxoacyl-[acyl-carrier-protein] reductase |
| 0.723 | A0A0I0MQ27 | Fimbrial protein (FimA) |
| 0.720 | A0A0I1VCS9 | HemG. Protoporphyrinogen IX dehydrogenase [quinone] |
| 0.719 | A0A0I0M9V1 | Lipoprotein |
| 0.717 | A0A0H9PY82 | Arginine ABC transporter periplasmic substrate-binding protein ArtJ |
| 0.716 | N/A | Hypothetical protein [*E. coli*] |
| 0.715 | A0A0H9RRA6 | ilvG Acetolactate synthase |
| 0.714 | A0A0I4E586 | Cytochrome c-type biogenesis protein; Haem chaperone required for the biogenesis of c-type cytochromes |
| 0.704 | A0A0I1Z9C6 | Hydroxyacylglutathione hydrolase |
| 0.704 | A0A0H7UZD9 | etp protein-tyrosine-phosphatase |
| 0.703 | A0A0H7L9I0 | RhaB. Rhamnulokinase. |

\*N/A refers to smaller peptides not found on the UniProt database, and the function is based on the top BLASTp hit

Furthermore, two different proteins with haem related functions with ipTM values ranging from 0.714 to 0.820 (NrfF, a possible subunit of a haem lyase and Cytochrome c-type biogenesis protein, a haem chaperone required for the biogenesis of c-type cytochromes respectively) (Table 1, Supplementary Table 4) suggesting Stv protein could be involved in *Shigella* virulence that might be important for successful pathogenesis, as haem is a critical nutrient/cofactor and the most abundant iron-containing compound in the host (38). The remaining majority of proteins with a high ipTM are mainly involved in metabolic functions, which may indicate the involvement of those protein in metabolic shift that occur during infection and intracellular growth (39).

## Discussion

We used historical and contemporary *Shigella* WGS collections to investigate factors driving its emergence as a successful pathogen. Historical isolates collections are a valuable resource to compare and contrast pathogen features to those in the modern isolates to pinpoint factors that have helped shape the pathogen trajectory not only in *Shigella* but also in other species such as *Klebsiella* spp. (18,20). Our novel approach, which we termed temporal GWAS (tGWAS), where genetic factors positively associated with time as a continuous variable in bacterial GWAS provides a novel tool to the bacterial GWAS repertoire. The results from our tGWAS analysis included known AMR genes (Supplementary Table 2) that was a positive and confirmatory result to validate the utility of tGWAS. Furthermore, we were able to identify genes for the *Shigella* T3SS that were carried on the well-known virulence plasmid (28,29) from our tGWAS analysis further confirming the validity of tGWAS as an effective method in identifying genetic factors associated with pathogenesis of *Shigella*. The recovery of known AMR and virulence factors also provide further confidence in the novel genes we discovered through tGWAS and demonstrates the potential utility of tGWAS in other pathogen datasets beyond *Shigella* as well.

From our tGWAS results, we focused on *stv*, the gene with the most associated k-mers but no functional annotation.The closest relative in sequence similarity searches to Stv was an *E. coli* protein annotated as an adhesin. However, we could not find any previous studies functionally validating this highlighting the limitations of annotation transfer. Further *in silico* analyses revealed Stv to contain a unique protein domain; now deposited in the Pfam database with accession PF21527. This domain was mainly located in the C-terminus of a variety of protein architectures with some architectures suggestive of a bacterial toxin (33,34). Expanding on our *in-silico* approach, we carried out an AlphaFold interaction screen looking at predicted protein-protein interactions using the *Shigella* proteome, resulted in high-confidence predictions of Stv as potentially interacting with fimbrial proteins and proteins with functions related to haem which may have potential roles in virulence. The interactions with fimbriae are particularly intriguing as *Shigella* is thought to canonically not have fimbriae due to their immunogenic properties to better hide themselves from a host immune response (17). The more successful lineages of Shigella are reported to have lost some of the *fim* operon genes responsible for type 1 fimbriae (40), while remnants have been observed in some lineages and the interactions of Stv with the Fim genes could be of an important factor during infection.

A potential role in virulence is consistent with the ecological evidence supporting the importance of *stv*. Specifically, that it is found associated with the epidemiologically successful *S. sonnei* Lineage 3, where the acquisition of pSTV predates the acquisition of the MDR conferring Tn*7*/Int2 cassette. Moreover, that pSTV has been maintained in bacterial populations despite being a small low-copy number plasmid with no AMR genes or addiction systems (41) suggests that it is likely contributing to the success (or public health visibility) of the pathogen (e.g. through increased virulence). Although we have narrowed down the potential interaction partners of Stv, laboratory studies are needed to confirm binding partners and further elucidate associated phenotypes.

In summary, our novel tGWAS approach is a powerful and generalisable tool for discovering novel proteins relevant to bacterial pathogens in an era of expanding WGS data (42). Using tGWAS we identified a novel Pfam domain whose ecological distribution and structural features point to a role in and in *Shigella’s* success as a bacterial pathogen.

## Materials and methods

### Whole genome sequence data and processing

Two publicly available *S. flexneri* whole genome sequenced datasets were used in this study where the first contained a collection of historical isolates from the Murray collection (n=45, isolated between 1917 – 1935) that can be accessed from ENA project number PM463261343GB (19). The second dataset consisted of a collection of modern isolates from a global study of *S. flexneri* (n=262, isolated between 1950 – 2011) (23). In addition to that, genome sequence data from NCTC1 (GCA_000953045.1) was also incorporated during the statistical analyses in the Murray isolates (13). *S. flexneri* 2a strain 301 (NC_004337.2) and its virulence plasmid (NC_004851.1 – pCP301) was used as a reference strain.

Three whole genome sequenced datasets were employed for analyses involving *S. sonnei*, where the first contained historical isolates from the Murray collection (n=22, collected from 1937 – 1954) accessible from the above ENA project. The second dataset originated from isolates from an investigation into the global population structure of *S. sonnei* (n=132) (30) along with supplementary isolates (n=24) from (12). The third dataset (n=127) consists of a subset of isolates representative of *S. sonnei* clade assignments from (43) and was used as the modern *S. sonnei* isolates in this study. *S. sonnei* 53G (HE616528.1) and its associated plasmids (H616529.1, H616530.1, H616531.1, H616532.1) were used as the reference strain.

Accession numbers and metadata for the strains used is listed in Supplementary Tables 1 and 3). Raw sequence data were adapter – and quality – trimmed using Trimmomatic v0.38 (44) and draft genome were assembled using Unicycler v0.4.7 (45) while annotations were carried out using Prokka v1.13.3 (46).

Phylogeny reconstruction and AMR and virulence determinants detection Trimmed FASTQ sequences were mapped against the relevant reference strains using Burrows – Wheeler Aligner (BWA) v0.5.9-r16 (47). The mapping files were then filtered and sorted using samtools v1.13 (48) while the duplicates were marked using Picard (49) and variant calling was completed with bcftools to generate a consensus genome sequence for each isolate (50). Each chromosome sequence was extracted and regions containing plasmid sequences, IS elements and repeat regions that were identified from the reference sequences were masked with a custom mask each for both *S. flexneri* and *S. sonnei*. Gubbins v2.3.4 was used to remove duplicate and low-quality sequences followed by SNP sites to obtain the core genome alignment (51,52) and phylogenetic tree was then inferred using RAxML-ng (53). Each of the phylogenetic trees were mid-point rooted and visualised using interactive Tree of Life (iTOL) v6.1.1(54).

Identification of genetic determinants attributable to AMR and virulence was performed using abricate v0.8.13 (55) using the NCBI AMRFinder Plus database and Virulence Factor Database (VFDB) (56,57) with a minimum identity threshold of 95.

### Temporal Genome Wide Association Study (tGWAS) and prioritisation strategy for GWAS hits

Paired end reads from *S. flexneri* isolates were mapped to the reference *S. flexneri* 2a strain 301 using BWA mem v07.17 (47) and Picard v2.23.1 was used to mark duplicates (49). VCF files generated after variant calling and subsequent filtering using Freebayes v1.3.2 (58) were merged and used as an input for the GWAS single nucleotide polymorphisms (SNPs) analysis. Small substrings of nucleotides of varying length of k from genome assemblies that were generated using fsm-lite v1.0 were used to generate the input for the GWAS kmer analysis. Gene_presence_absence.Rtab file resulting from pan genome calculations using Roary v1.007002 (59) was used for the primary clustering of Clusters of Orthologous Genes (COGs) and cross referencing of significant COGs with a secondary COG GWAS analysis was carried out using the gene_presence_absence.Rtab file generated from pangenome calculations from Panaroo v1.2.10 (60). GWAS was conducted using Pyseer v1.3.6 (61) using the appropriate input files mentioned above. For the specific temporal GWAS, time in years, was used as the continuous phenotype with Pyseer. All analyses were supplemented with phylogenetic distances from the mid-point rooted phylogeny and a covariate file of assigned phylogroups to account for population structure when using the linear mixed model (LMM) of Pyseer. SnpEff v4.3.1 (62) was used to further explore and predict the functional effects of annotated variants of SNP hits generated from the GWAS. A custom-made decision tree (Supplementary Figure 5) was developed to prioritise the hits from the three GWAS analysis types and to account for each genetic feature type.

### Protein tree construction

Amino acid sequence of Stv was utilised in a BLASTp search against the clustered nr database. Sequences belonging to the cluster identified in this search (63) were extracted and aligned with the Stv sequence using muscle v5.1 (64) after which RAxML-ng v8.2.9 (53) was used to construct a phylogeny based on the alignment with 100 bootstraps.

### Identification and confirmation of pSTV

Both *in silico* and *in vitro* methods were used to confirm the genomic context and copy number of *stv* in the genome of the corresponding clinical isolate SRR1364216 (8). Paired end reads were mapped back onto the assembly via Burrows-Wheeler Aligner v 0.5.9-r16 (47) and filtered and sorted using samtools v1.13 (48) to determine the copy number difference for *stv* and the chromosome. Coverage was calculated using samtools depth v1.13 and average was calculated for both the chromosomal contig and the smaller *stv* containing contig. Full reconstruction of the pSTV sequence (Genbank accession number: OP113953) was made using BLAST and Artemis Comparison Tool (ACT) comparison of the stv-containing contiguous sequence in multiple isolates while the sequence was confirmed via PCR analysis of genomic DNA extracted from ERR1364216 (Supplementary Figure 2)

### Domain analysis and AlphaFold screening

Stv homologues were identified using iterative HMMer searches with the hmmsearch program against the UniProt reference proteome sequences using a per-sequence and per-domain threshold of 27 bits. The resulting alignment was submitted to Pfam for inclusion and assigned accession number PF21527.

Interactions between Stv and the proteome of the Shigella strain 627346 were predicted using AlphaPullDown v1.0.4 (65). Of the 5,239 candidate complexes, 4,859 predictions were successfully modelled with AlphaFold2 (66), while 380 failed due to size constraints and were subsequently modelled with AlphaFold3 (version AlphaFold-beta-20231127) (67). All modelling was performed on Tesla V100 32GB GPUs. Functional annotations for the proteins involved in significant interactions (ipTM > 0.7) were downloaded from the UniProtKB website (database version 2025_03) (68) using the query “(taxonomy_id:620)”.

## Acknowledgements

Rebecca J Bennett is funded by a Biotechnology and Biological Sciences Research Council Doctoral Training Partnership studentship (BB/M011186/1). KSB was supported by funding from the MRC (MR/R020787/1) and BBSRC (BB/V009184/1). KSB is also affiliated to the National Institute for Health Research Health Protection Research Unit (NIHR HPRU) in Gastrointestinal Infections at University of Liverpool in partnership with the United Kingdom Health Security Agency, in collaboration with University of Warwick. The views expressed are those of the author(s) and not necessarily those of the NHS, the NIHR, the Department of Health and Social Care or UKHSA. The authors are grateful to John Lees, Sarah Alexander, and Anna Auer-Fowler helpful discussions of work in progress.

## Notes

### Competing Interest Statement

Tim Blower is a current employee of New England Biolabs, but was not during the time of this work.

### Summary of Updates

The manuscript has been condensed to create room for new Stv protein investigation results in collaboration with EBI including model, domain architecture exploration, and in silico pull down assays.

## References

1. Hendrick J, Raqib R, Noor Z, Faruque ASG, Haque R, Petri WA. Shigellosis. Lancet. 2025 Oct 4;406(10511):1508–19.

2. Scott TA, Baker KS, Trotter C, Jenkins C, Mostowy S, Hawkey J, et al. Shigella sonnei: epidemiology, evolution, pathogenesis, resistance and host interactions. Nat Rev Microbiol. 2024 Nov 27;

3. Kotloff KL, Riddle MS, Platts-Mills JA, Pavlinac P, Zaidi AKM. Shigellosis. Lancet. 2018 Feb 24;391(10122):801–12.

4. Livio S, Strockbine NA, Panchalingam S, Tennant SM, Barry EM, Marohn ME, et al. Shigella isolates from the global enteric multicenter study inform vaccine development. Clin Infect Dis. 2014 Oct;59(7):933–41.

5. Kasumba IN, Badji H, Powell H, Hossain MJ, Omore R, Sow SO, et al. Shigella in Africa: New insights from the Vaccine Impact on diarrhea in Africa (VIDA) study. Clin Infect Dis. 2023 Apr 19;76(76 Suppl1):S66–76.

6. Anderson JD 4th, Bagamian KH, Muhib F, Amaya MP, Laytner LA, Wierzba T, et al. Burden of enterotoxigenic Escherichia coli and shigella non-fatal diarrhoeal infections in 79 low-income and lower middle-income countries: a modelling analysis. Lancet Glob Health. 2019 Mar;7(3):e321–30.

7. Khalil IA, Troeger C, Blacker BF, Rao PC, Brown A, Atherly DE, et al. Morbidity and mortality due to shigella and enterotoxigenic Escherichia coli diarrhoea: the Global Burden of Disease Study 1990-2016. Lancet Infect Dis. 2018 Nov;18(11):1229–40.

8. Baker KS, Dallman TJ, Field N, Childs T, Mitchell H, Day M, et al. Horizontal antimicrobial resistance transfer drives epidemics of multiple Shigella species. Nat Commun. 2018 Apr 13;9(1):1462.

9. Baker KS, Dallman TJ, Behar A, Weill F-X, Gouali M, Sobel J, et al. Travel- and community-based transmission of multidrug-resistant Shigella sonnei lineage among international orthodox Jewish communities. Emerg Infect Dis. 2016 Sept;22(9):1545–53.

10. Mason LCE, Greig DR, Cowley LA, Partridge SR, Martinez E, Blackwell GA, et al. The evolution and international spread of extensively drug resistant Shigella sonnei. Nat Commun. 2023 Apr 8;14(1):1983.

11. Bardsley M, Jenkins C, Mitchell HD, Mikhail AFW, Baker KS, Foster K, et al. Persistent transmission of shigellosis in England is associated with a recently emerged multidrug-resistant strain of Shigella sonnei. J Clin Microbiol. 2020 Mar 25;58(4):e01692–19.

12. Baker KS, Dallman TJ, Field N, Childs T, Mitchell H, Day M, et al. Genomic epidemiology of Shigella in the United Kingdom shows transmission of pathogen sublineages and determinants of antimicrobial resistance. Sci Rep. 2018 May 9;8(1):7389.

13. Baker KS, Mather AE, McGregor H, Coupland P, Langridge GC, Day M, et al. The extant World War 1 dysentery bacillus NCTC1: a genomic analysis. Lancet. 2014 Nov 8;384(9955):1691–7.

14. Sati H, Carrara E, Savoldi A, Hansen P, Garlasco J, Campagnaro E, et al. The WHO Bacterial Priority Pathogens List 2024: a prioritisation study to guide research, development, and public health strategies against antimicrobial resistance. Lancet Infect Dis. 2025 Sept;25(9):1033–43.

15. Baker KS, Dallman TJ, Ashton PM, Day M, Hughes G, Crook PD, et al. Intercontinental dissemination of azithromycin-resistant shigellosis through sexual transmission: a cross-sectional study. Lancet Infect Dis. 2015 Aug;15(8):913–21.

16. De Silva PM, Bennett RJ, Kuhn L, Ngondo P, Debande L, Njamkepo E, et al. Escherichia coli killing by epidemiologically successful sublineages of Shigella sonnei is mediated by colicins. EBioMedicine. 2023 Nov;97(104822):104822.

17. Miles SL, Santillo D, Painter H, Wright K, Torraca V, López-Jiménez AT, et al. Enhanced virulence and stress tolerance are signatures of epidemiologically successful Shigella sonnei. Nat Commun. 2025 Oct 9;16(1):9005.

18. Bennett RJ, Baker KS. Looking backward to move forward: The utility of sequencing historical bacterial genomes. J Clin Microbiol [Internet]. 2019 Aug;57(8). Available from: 10.1128/JCM.00100-19

19. Baker KS, Burnett E, McGregor H, Deheer-Graham A, Boinett C, Langridge GC, et al. The Murray collection of pre-antibiotic era Enterobacteriacae: a unique research resource. Genome Med. 2015 Sept 28;7(1):97.

20. Wand ME, Baker KS, Benthall G, McGregor H, McCowen JWI, Deheer-Graham A, et al. Characterization of pre-antibiotic era Klebsiella pneumoniae isolates with respect to antibiotic/disinfectant susceptibility and virulence in Galleria mellonella. Antimicrob Agents Chemother. 2015 July;59(7):3966–72.

21. Hughes VM, Datta N. Conjugative plasmids in bacteria of the ‘pre-antibiotic’ era. Nature. 1983 Apr 21;302(5910):725–6.

22. Cazares A, Figueroa W, Cazares D, Lima L, Turnbull JD, McGregor H, et al. Pre- and postantibiotic epoch: The historical spread of antimicrobial resistance. Science. 2025 Dec 4;390(6777):eadr1522.

23. Connor TR, Barker CR, Baker KS, Weill F-X, Talukder KA, Smith AM, et al. Species-wide whole genome sequencing reveals historical global spread and recent local persistence in Shigella flexneri. Elife. 2015 Aug 4;4:e07335.

24. Njamkepo E, Fawal N, Tran-Dien A, Hawkey J, Strockbine N, Jenkins C, et al. Global phylogeography and evolutionary history of Shigella dysenteriae type 1. Nat Microbiol. 2016 Mar 21;1(4):16027.

25. Hammarlöf DL, Kröger C, Owen SV, Canals R, Lacharme-Lora L, Wenner N, et al. Role of a single noncoding nucleotide in the evolution of an epidemic African clade of Salmonella. Proc Natl Acad Sci U S A. 2018 Mar 13;115(11):E2614–23.

26. Horiyama T, Nishino K. AcrB, AcrD, and MdtABC multidrug efflux systems are involved in enterobactin export in Escherichia coli. PLoS One. 2014 Sept 26;9(9):e108642.

27. Møller Tsb, Overgaard M, Nielsen SS, Bortolaia V, Sommer MOA, Guardabassi L, et al. Relation between tetR and tetA expression in tetracycline resistant Escherichia coli. BMC Microbiol. 2016 Mar 12;16(1):39.

28. Bajunaid W, Haidar-Ahmad N, Kottarampatel AH, Ourida Manigat F, Silué N, Tchagang CF, et al. The T3SS of Shigella: Expression, structure, function, and role in vacuole escape. Microorganisms. 2020 Dec 5;8(12):1933.

29. Ogawa M, Handa Y, Ashida H, Suzuki M, Sasakawa C. The versatility of Shigella effectors. Nat Rev Microbiol. 2008 Jan;6(1):11–6.

30. Holt KE, Baker S, Weill F-X, Holmes EC, Kitchen A, Yu J, et al. Shigella sonnei genome sequencing and phylogenetic analysis indicate recent global dissemination from Europe. Nat Genet. 2012 Sept;44(9):1056–9.

31. Caryl JA, Smith MCA, Thomas CD. Reconstitution of a staphylococcal plasmid-protein relaxation complex in vitro. J Bacteriol. 2004 June;186(11):3374–83.

32. Castagnoli L, Scarpa M, Kokkinidis M, Banner DW, Tsernoglou D, Cesareni G. Genetic and structural analysis of the ColE1 Rop (Rom) protein. EMBO J. 1989 Feb;8(2):621–9.

33. Busby JN, Panjikar S, Landsberg MJ, Hurst MRH, Lott JS. The BC component of ABC toxins is an RHS-repeat-containing protein encapsulation device. Nature. 2013 Sept 26;501(7468):547–50.

34. Ali J, Yu M, Sung L-K, Cheung Y-W, Lai E-M. A glycine zipper motif is required for the translocation of a T6SS toxic effector into target cells. EMBO Rep. 2023 June 5;24(6):e56849.

35. Julien O, Wells JA. Caspases and their substrates. Cell Death Differ. 2017 Aug;24(8):1380–9.

36. Asplund-Samuelsson J, Bergman B, Larsson J. Prokaryotic caspase homologs: phylogenetic patterns and functional characteristics reveal considerable diversity. PLoS One. 2012 Nov 19;7(11):e49888.

37. Shen A, Lupardus PJ, Albrow VE, Guzzetta A, Powers JC, Garcia KC, et al. Mechanistic and structural insights into the proteolytic activation of Vibrio cholerae MARTX toxin. Nat Chem Biol. 2009 July;5(7):469–78.

38. Choby JE, Skaar EP. Heme synthesis and acquisition in bacterial pathogens. J Mol Biol. 2016 Aug 28;428(17):3408–28.

39. Kentner D, Martano G, Callon M, Chiquet P, Brodmann M, Burton O, et al. Shigella reroutes host cell central metabolism to obtain high-flux nutrient supply for vigorous intracellular growth. Proc Natl Acad Sci U S A. 2014 July 8;111(27):9929–34.

40. Bravo V, Puhar A, Sansonetti P, Parsot C, Toro CS. Distinct mutations led to inactivation of type 1 fimbriae expression in Shigella spp. PLoS One. 2015 Mar 26;10(3):e0121785.

41. Castañeda-Barba S, Top EM, Stalder T. Plasmids, a molecular cornerstone of antimicrobial resistance in the One Health era. Nat Rev Microbiol. 2024 Jan;22(1):18–32.

42. Perez-Sepulveda BM, Heavens D, Pulford CV, Predeus AV, Low R, Webster H, et al. An accessible, efficient and global approach for the large-scale sequencing of bacterial genomes. Genome Biol. 2021 Dec 21;22(1):349.

43. Hawkey J, Paranagama K, Baker KS, Bengtsson RJ, Weill F-X, Thomson NR, et al. Global population structure and genotyping framework for genomic surveillance of the major dysentery pathogen, Shigella sonnei. Nat Commun. 2021 May 11;12(1):2684.

44. Bolger AM, Lohse M, Usadel B. Trimmomatic: a flexible trimmer for Illumina sequence data. Bioinformatics. 2014 Aug 1;30(15):2114–20.

45. Wick RR, Judd LM, Gorrie CL, Holt KE. Unicycler: Resolving bacterial genome assemblies from short and long sequencing reads. PLoS Comput Biol. 2017 June;13(6):e1005595.

46. Seemann T. Prokka: rapid prokaryotic genome annotation. Bioinformatics. 2014 July 15;30(14):2068–9.

47. Li H, Durbin R. Fast and accurate short read alignment with Burrows-Wheeler transform. Bioinformatics. 2009 July 15;25(14):1754–60.

48. Li H, Handsaker B, Wysoker A, Fennell T, Ruan J, Homer N, et al. The Sequence Alignment/Map format and SAMtools. Bioinformatics. 2009 Aug 15;25(16):2078–9.

49. Broad Institute. Picard Toolkit. https://broadinstitute.github.io/picard/. 2019.

50. Danecek P, McCarthy SA. BCFtools/csq: haplotype-aware variant consequences. Bioinformatics. 2017 July 1;33(13):2037–9.

51. Page AJ, Taylor B, Delaney AJ, Soares J, Seemann T, Keane JA, et al. SNP-sites: rapid efficient extraction of SNPs from multi-FASTA alignments. Microb Genom. 2016 Apr;2(4):e000056.

52. Croucher NJ, Page AJ, Connor TR, Delaney AJ, Keane JA, Bentley SD, et al. Rapid phylogenetic analysis of large samples of recombinant bacterial whole genome sequences using Gubbins. Nucleic Acids Res. 2015 Feb 18;43(3):e15.

53. Kozlov AM, Darriba D, Flouri T, Morel B, Stamatakis A. RAxML-NG: a fast, scalable and user-friendly tool for maximum likelihood phylogenetic inference. Bioinformatics. 2019 Nov 1;35(21):4453–5.

54. Letunic I, Bork P. Interactive Tree of Life (iTOL) v6: recent updates to the phylogenetic tree display and annotation tool. Nucleic Acids Res. 2024 July 5;52(W1):W78–82.

55. Seemann T. Abricate, Github. https://github.com/tseemann/abricate.

56. Feldgarden M, Brover V, Haft DH, Prasad AB, Slotta DJ, Tolstoy I, et al. Validating the AMRFinder tool and resistance gene database by using antimicrobial resistance genotype-phenotype correlations in a collection of isolates. Antimicrob Agents Chemother. 2019 Nov;63(11).

57. Chen L, Zheng D, Liu B, Yang J, Jin Q. VFDB 2016: hierarchical and refined dataset for big data analysis--10 years on. Nucleic Acids Res. 2016 Jan 4;44(D1):D694–7.

58. Garrison E, Marth G. Haplotype-based variant detection from short-read sequencing [Internet]. arXiv [q-bio.GN]. 2012. Available from: http://arxiv.org/abs/1207.3907

59. Page AJ, Cummins CA, Hunt M, Wong VK, Reuter S, Holden MTG, et al. Roary: rapid large-scale prokaryote pan genome analysis. Bioinformatics. 2015 Nov 15;31(22):3691–3.

60. Tonkin-Hill G, MacAlasdair N, Ruis C, Weimann A, Horesh G, Lees JA, et al. Producing polished prokaryotic pangenomes with the Panaroo pipeline. Genome Biol. 2020 July 22;21(1):180.

61. Lees JA, Galardini M, Bentley SD, Weiser JN, Corander J. Pyseer: A comprehensive tool for microbial pangenome-wide association studies. Bioinformatics. 2018 Dec 15;34(24):4310–2.

62. Cingolani P, Platts A, Wang LL, Coon M, Nguyen T, Wang L, et al. A program for annotating and predicting the effects of single nucleotide polymorphisms, SnpEff: SNPs in the genome of Drosophila melanogaster strain w1118; iso-2; iso-3. Fly (Austin). 2012 Apr;6(2):80–92.

63. Steinegger M, Söding J. MMseqs2 enables sensitive protein sequence searching for the analysis of massive data sets. Nat Biotechnol. 2017 Nov;35(11):1026–8.

64. Edgar RC. MUSCLE: multiple sequence alignment with high accuracy and high throughput. Nucleic Acids Res. 2004 Mar 19;32(5):1792–7.

65. Yu D, Chojnowski G, Rosenthal M, Kosinski J. AlphaPulldown-a python package for protein-protein interaction screens using AlphaFold-Multimer. Bioinformatics. 2023 Jan 1;39(1):btac749.

66. Jumper J, Evans R, Pritzel A, Green T, Figurnov M, Ronneberger O, et al. Highly accurate protein structure prediction with AlphaFold. Nature. 2021 Aug;596(7873):583–9.

67. Abramson J, Adler J, Dunger J, Evans R, Green T, Pritzel A, et al. Accurate structure prediction of biomolecular interactions with AlphaFold 3. Nature. 2024 June;630(8016):493–500.

68. Ahmad S, Jose da Costa Gonzales L, Bowler-Barnett EH, Rice DL, Kim M, Wijerathne S, et al. The UniProt website API: facilitating programmatic access to protein knowledge. Nucleic Acids Res. 2025 July 7;53(W1):W547–53.

