## Supplementary Figures for "Temporal GWAS identifies a widely distributed novel protein Stv contributing to pathogen success in *Shigella* spp"

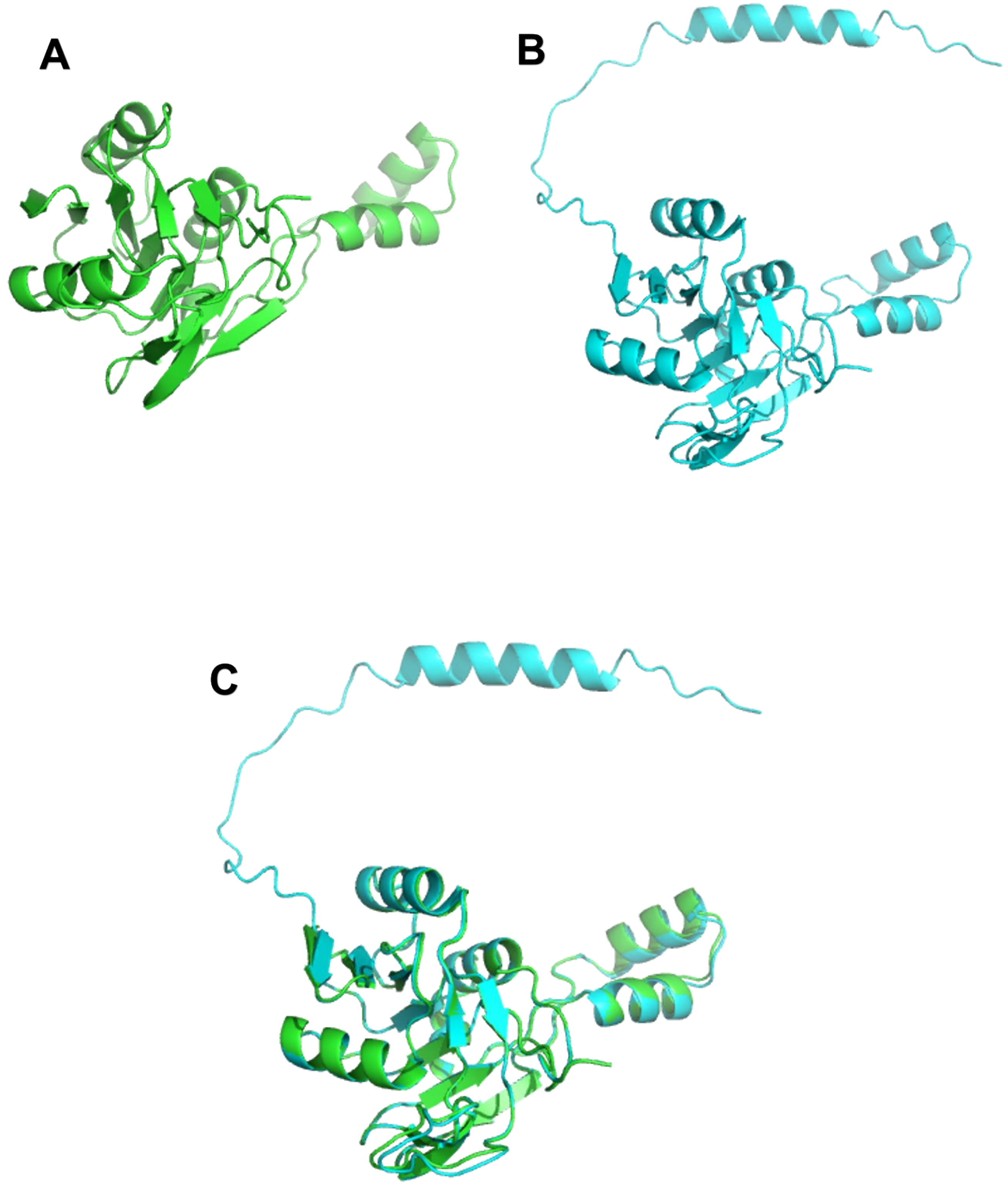


Supplementary Figure 1. AlphaFold 2 modelling of (A) *E. coli* adhesin in green and (B) *stv* in cyan and the (C) overlay of the two showing the structural similarity and the c-terminal loop that has no matching regions in the *E. coli* adhesin

**A**

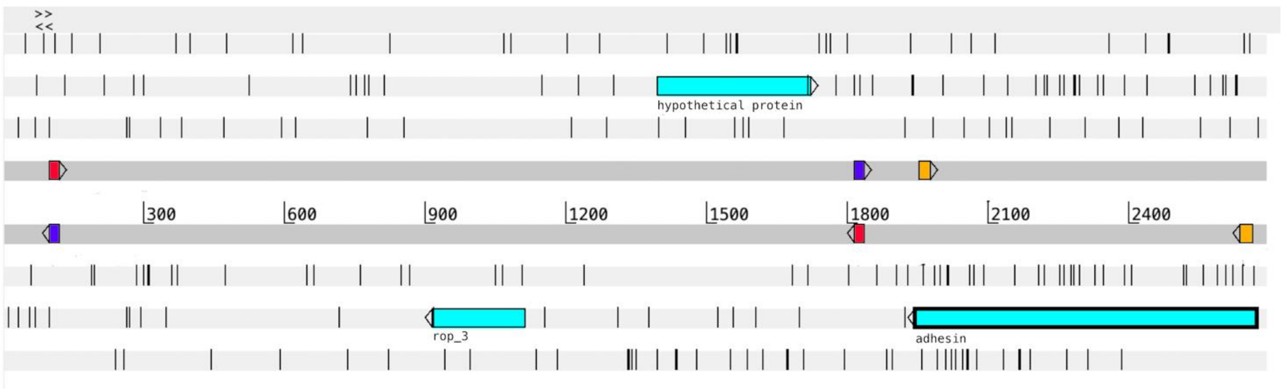


| Primers | Sequence | Expected product size |
|---|---|---|
| Outward_F<br>Outward_R | ggatctcaagaagaaccttt<br>tgaaaacgatcctgacgcat | 993bp |
| Inward_F<br>Inward_R | aaaggttcttcttgagatcc<br>atgcgtcaggatcgttttca | 1737bp |
| Internal_F<br>Internal_R | actcaattcgaatctcataacaaa<br>cctctctatttactcggcaagc | 710bp |

**B**

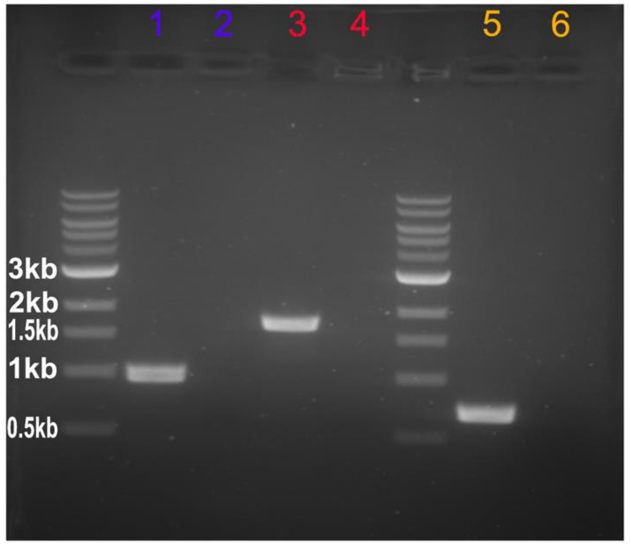


1 - Outward region
2 - No template control

3 - Inward region
4 - No template control

5 - Internal region
6 - No template control

**C**

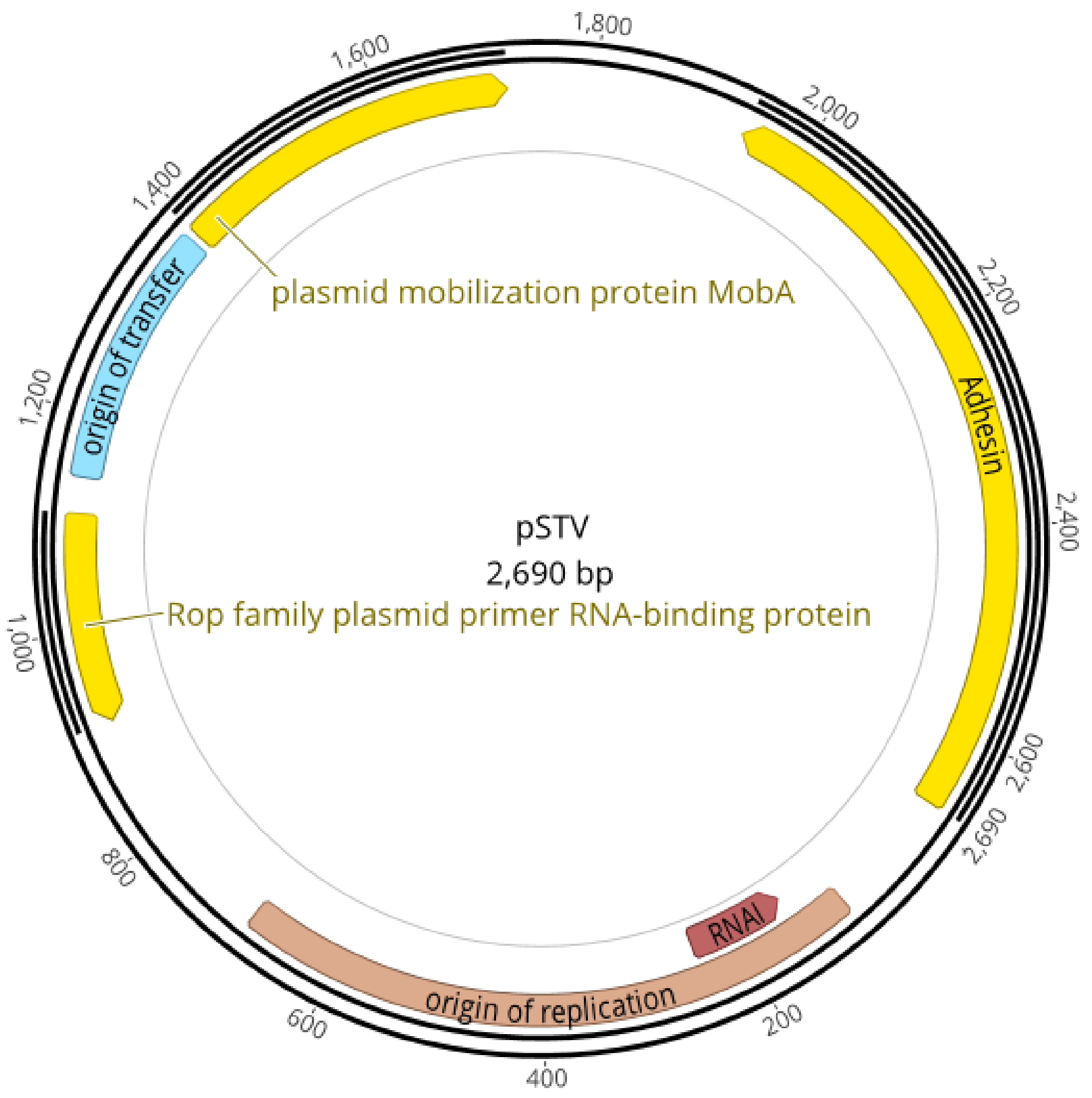


Supplementary Figure 2. Confirmation of adhesin Stv on pStv by PCR. (A) A linear map of the contiguous sequence containing adehsin extracted from ERR1364216 is shown with gene annotations. Primer sequences designed to amplify the plasmid interior and exterior to the contig boundary as well as internal region of the adhesin gene are shown in coloured blocks, with the sequences and the expected product sizes from PCR in the table below. (B) Gel image of three PCR reactions along with their relevant no template controls for the respective primer pairs were conducted and demonstrated, through their being the expected size, that the sequence is complete. Based on the evidence from the PCR reactions and genome coverage information, overlapping contiguous sequences were used to manually edit a final circular pSTV sequence, which is deposited in GenBank under accession number: OP113953. (C) Plasmid map of pSTV with the features indicated on the map

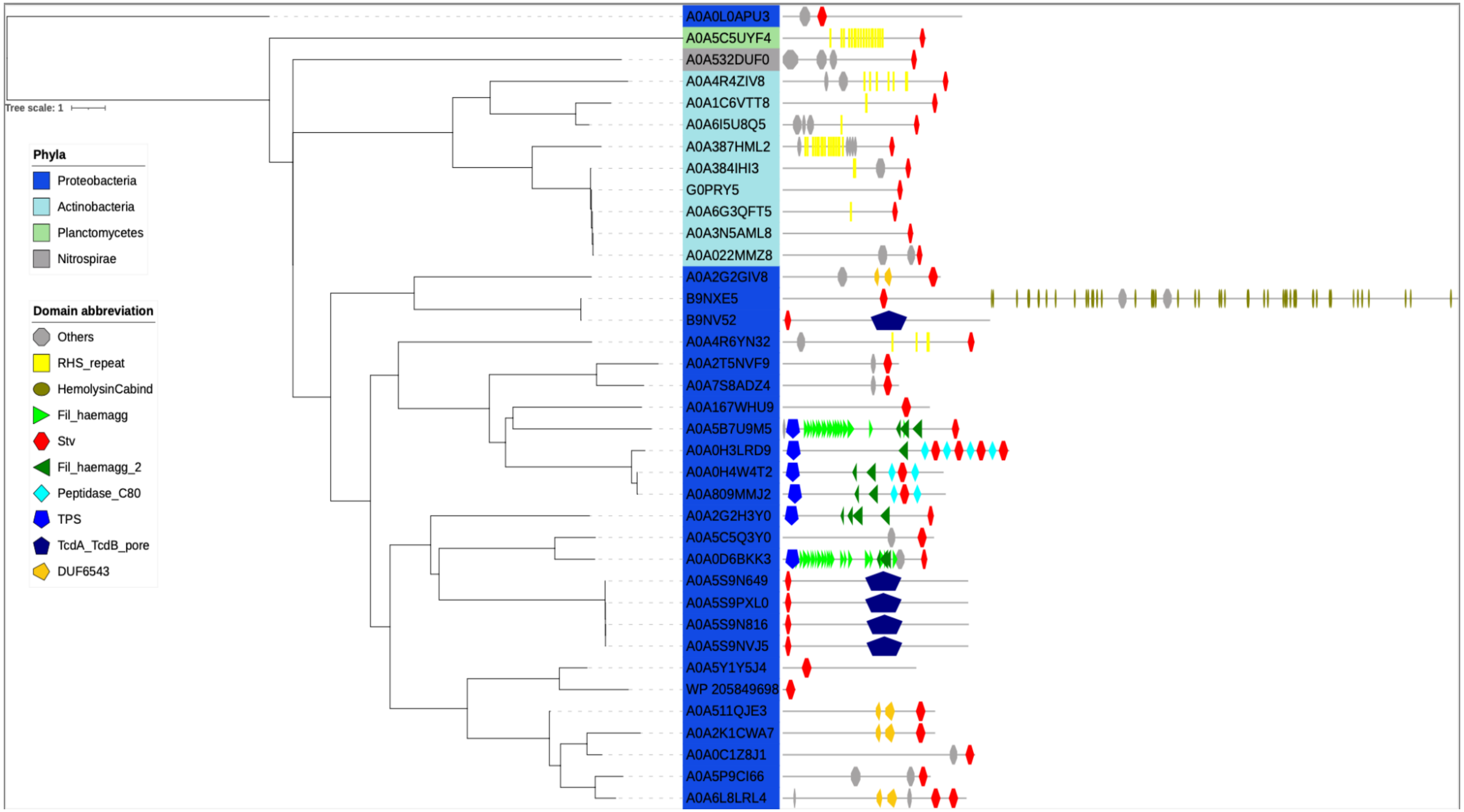


Supplementary Figure 3. Domain architecture of selection of Stv domain containing proteins considering their taxonomic relation: The phylogenetic tree is generated from 16S rRNA sequences of the genera of the selection of Stv domain containing proteins. The illustrated domains are found in the Pfam database with the following accession numbers: RHS_repeat:PF05593, HemolysinCabind:PF00353, Fil_haemagg:PF05594, Fil_haemagg_2:PF13332, Stv:PF21527, Peptidase_C80:PF11713, TPS:PF05860, TcdA_TcdB_pore:PF12920, DUF6543:PF20178.

| No: | Chain | Z | rmsd | lali | nres | %id | PDB | Description |
|---|---|---|---|---|---|---|---|---|
| 1: | 3gff-A | 4.0 | 3.6 | 83 | 316 | 10 | PDB | MOLECULE: IROE-LIKE SERINE HYDROLASE; |
| 2: | 7d78-D | 3.8 | 3.7 | 85 | 297 | 12 | PDB | MOLECULE: DLTD DOMAIN-CONTAINING PROTEIN; |
| 3: | 5x6s-A | 3.7 | 3.5 | 82 | 274 | 9 | PDB | MOLECULE: ACETYLXYLAN ESTERASE A; |
| 4: | 2xt6-A | 3.6 | 4.8 | 75 | 1055 | 9 | PDB | MOLECULE: 2-OXOGLUTARATE DECARBOXYLASE; |
| 5: | 4hxg-F | 3.5 | 3.9 | 82 | 614 | 10 | PDB | MOLECULE: PUTATIVE UNCHARACTERIZED PROTEIN PH0594; |
| 6: | 6mou-A | 3.5 | 4.5 | 81 | 359 | 10 | PDB | MOLECULE: ISOAMYLASE N-TERMINAL DOMAIN PROTEIN; |
| 7: | 2o2g-A | 3.5 | 4.1 | 84 | 216 | 11 | PDB | MOLECULE: DIENELACTONE HYDROLASE; |
| 8: | 6wym-A | 3.4 | 3.2 | 76 | 291 | 7 | PDB | MOLECULE: POSSIBLE HYDROLASE; |
| 9: | 4x00-A | 3.3 | 3.0 | 75 | 273 | 8 | PDB | MOLECULE: PUTATIVE HYDROLASE; |
| 10: | 7dwc-A | 3.3 | 3.0 | 81 | 264 | 12 | PDB | MOLECULE: XYLANASE; |
| 11: | 7jsr-A | 3.3 | 2.9 | 78 | 1496 | 4 | PDB | MOLECULE: NAD-SPECIFIC GLUTAMATE DEHYDROGENASE; |
| 12: | 5g59-A | 3.3 | 3.4 | 80 | 274 | 9 | PDB | MOLECULE: ESTERASE; |
| 13: | 6gi5-B | 3.3 | 3.4 | 80 | 272 | 13 | PDB | MOLECULE: FERRIC ENTEROBACTIN ESTERASE; |
| 14: | 2q0x-A | 3.2 | 3.8 | 78 | 294 | 8 | PDB | MOLECULE: UNCHARACTERIZED PROTEIN; |
| 15: | 3f67-A | 3.1 | 3.6 | 80 | 240 | 5 | PDB | MOLECULE: PUTATIVE DIENELACTONE HYDROLASE; |
| 16: | 4ao8-A | 3.1 | 3.3 | 79 | 233 | 10 | PDB | MOLECULE: ESTERASE; |
| 17: | 3pfo-A | 3.1 | 4.6 | 80 | 426 | 4 | PDB | MOLECULE: PUTATIVE ACETYLORNITHINE DEACETYLASE; |
| 18: | 5gad-i | 3.0 | 3.9 | 87 | 450 | 10 | PDB | MOLECULE: ESRP 4.5S RNA; |
| 19: | 6gup-A | 3.0 | 3.9 | 83 | 340 | 2 | PDB | MOLECULE: SIDEROPHORE BIOSYNTHESIS LIPASE/ESTERASE, PUTATIV |
| 20: | 6khm-K | 2.9 | 3.0 | 79 | 312 | 9 | PDB | MOLECULE: HYDROLASE, ALPHA/BETA DOMAIN PROTEIN; |
| 21: | 4w9r-B | 2.9 | 4.2 | 81 | 361 | 7 | PDB | MOLECULE: UNCHARACTERIZED PROTEIN; |
| 22: | 6ica-C | 2.9 | 4.6 | 81 | 378 | 6 | PDB | MOLECULE: AMINOPEPTIDASE; |
| 23: | 1r88-A | 2.9 | 3.6 | 78 | 267 | 9 | PDB | MOLECULE: MPT51/MPB51 ANTIGEN; |
| 24: | 2fx5-A | 2.8 | 3.3 | 77 | 258 | 3 | PDB | MOLECULE: LIPASE; |
| 25: | 3mga-A | 2.8 | 3.5 | 83 | 397 | 8 | PDB | MOLECULE: ENTEROCHELIN ESTERASE; |
| 26: | 6z69-B | 2.8 | 3.7 | 77 | 364 | 5 | PDB | MOLECULE: ACETYL ESTERASE/LIPASE; |
| 27: | 7wab-A | 2.8 | 3.3 | 77 | 484 | 6 | PDB | MOLECULE: COMPASS (COMPLEX PROTEINS ASSOCIATED WITH SET1P) |

Supplementary Figure 4. The top DALI matches for Stv AlphaFold model (low Z-scores) are shown above. None of the matches has a high DALI score indicative of a homologous relationship.

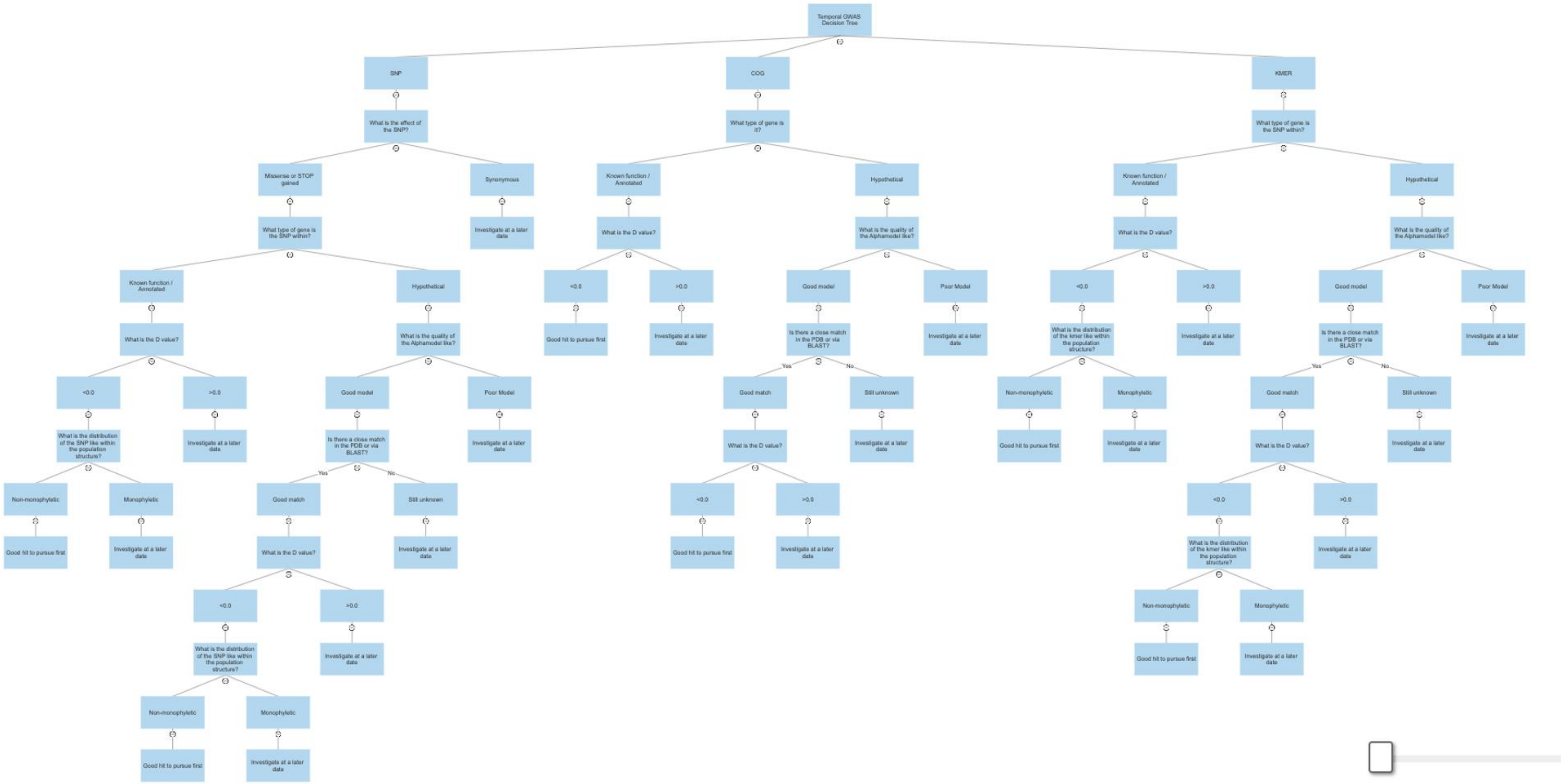


Supplementary Figure 5. Graphical representation of the decision trees allowing prioritisation of key tGWAS feature types – SNPs, kmers and COGs
